# MISSTE: a multiscale integrative spatial simulator for understanding the mechanisms underlying tissue ecosystems

**DOI:** 10.64898/2026.04.14.718434

**Authors:** Zhaoqian Su, Shanye Yin, Yinghao Wu

**Author notes:** Corresponding authors: Yinghao Wu.

## Abstract

Multiscale tissue ecosystems are governed by coupled intracellular decision-making, cell-cell interactions, and spatially structured microenvironmental signals, yet these scales are often studied separately. Here we present MISSTE, a modular framework that integrates Boolean intracellular state logic, agent-based modeling, and partial differential equation fields within a unified spatial simulation architecture. As a proof of concept, we applied MISSTE to CAR-T therapy in a solid tumor microenvironment. The model recapitulated emergent features of CAR-T behavior, including limited tumor penetration, stromal suppression, localized cytokine remodeling, hypoxia-associated constraint, and progressive functional exhaustion. Comparison of baseline and optimized conditions showed that coordinated enhancement of interaction range, migration, and cytotoxic function improved immune persistence and partial tumor control. Systematic parameter scans further identified effective immune-tumor contact as a stronger determinant of outcome than killing strength alone, highlighting spatial access as the dominant bottleneck. Guided by these results, we designed sequential intervention strategies and found that time-ordered enhancement of infiltration, killing, and late functional protection outperformed a static optimized regime. Together, these results establish MISSTE as a generalizable multiscale methodology for dissecting tissue ecosystems and for generating mechanistically grounded strategies for engineered cellular therapy design.

## 1. Introduction

The dynamics of multicellular organisms, such as tissue ecosystems, are governed by phenomena that span the molecular, cellular, and macroscopic scales [1]. At the microscale, intracellular signaling programs determine how individual cells sense and adapt to their environment [2]. At the mesoscale, direct cell-cell interactions drive collective behaviors such as immune activation, competition, and targeted suppression [3]. Simultaneously at the macroscale, diffusible microenvironmental factors regulate these localized events over broad spatial and temporal dimensions [4]. Because these biological layers are tightly coupled through continuous feedback, population-level outcomes cannot be predicted by isolating any single scale. For example, bulk tissue sequencing [5–9] captures macroscopic molecular signatures but obscures the physical cell-to-cell contacts and local micro-niches that drive immune exhaustion. On the other hand, advanced spatial transcriptomics [10–15] provides powerful architectural snapshots but lacks the temporal resolution required to capture the continuous, dynamic feedback between migrating cells and diffusing cytokine fields. Therefore, integrative frameworks are increasingly required to connect intracellular decision logic, multicellular interactions, and spatial tissue organization.

Chimeric antigen receptor (CAR) T-cell therapy [16–18] serves as an ideal test system for such a framework, presenting a profound intersection of methodological complexity and urgent biological relevance. Biologically, while CAR-T therapies have revolutionized the treatment of hematological malignancies, their efficacy in solid tumors is severely limited by the highly immunosuppressive tumor microenvironment (TME), where localized hypoxia, stromal barriers, and inhibitory signals rapidly drive T-cell exhaustion [19]. Methodologically, the dynamics of CAR-T therapy perfectly encapsulate the need for a multiscale approach. The system is driven by engineered intracellular logic (CAR signaling and state transitions), governed by direct mesoscale cellular interactions (immunological synapse formation and targeted cytotoxicity), and continuously modulated by macroscale tissue fields (the diffusion of inflammatory cytokines and suppressive gradients). Consequently, modeling CAR-T therapy not only rigorously validates the computational architecture but also provides a vital translational tool for dissecting immune failure and optimizing therapeutic design.

Computational modeling is increasingly used to decode tissue ecosystems [20–22], including CAR-T pharmacodynamics [23, 24]. Spatially averaged ordinary differential equations (ODEs) inherently cannot capture the profound spatial heterogeneity of the TME, such as physical exclusion and localized hypoxic gradients [25, 26]. Conversely, existing spatial models that utilize agent-based [27] or cellular automaton [28] frameworks successfully capture local interactions but frequently abstract away the intracellular signaling networks responsible for complex behaviors like antigen-induced exhaustion. Although fully coupled multiscale models have been proposed to bridge this gap, challenge still remains for a framework that simultaneously links intracellular signaling, cell-resolved interactions, and tissue-scale microenvironmental dynamics in a mechanistically interpretable and computationally scalable manner.

To address this challenge, we developed MISSTE (Multiscale Integrative Spatial Simulator for understanding the mechanisms underlying Tissue Ecosystems), a modular computational framework that integrates partial differential equations (PDEs) for continuum microenvironmental gradients, an agent-based model (ABM) for spatially explicit cell-cell interactions, and Boolean state networks for intracellular decision-making. We used MISSTE to model CAR-T therapy in a solid tumor microenvironment as a proof-of-concept system in which receptor-driven activation, spatially constrained immune-tumor contact, and suppressive tissue ecology interact across scales. In this setting, MISSTE recapitulated emergent features of CAR-T dynamics, including limited tumor penetration, localized cytokine accumulation, and antigen-dependent dysfunction/exhaustion. The simulations further suggested that therapeutic failure can arise not only from insufficient killing capacity, but from the coupled effects of spatial exclusion, stromal suppression, and metabolic constraint within the tumor microenvironment. By systematically scanning model parameters and intervention rules, we show that the framework can also be used to identify candidate strategies that improve immune persistence and tumor control. Because intracellular logic, cell behaviors, and microenvironmental interactions are encoded as interchangeable rule-based modules, MISSTE is readily extensible to other tissue ecosystems and engineered cell systems. More broadly, it provides a generalizable computational foundation for integrating spatial single-cell information, interrogating multicellular mechanisms, and supporting the rational design of next-generation cellular therapies.

## 2. Model and Methods

### 2.1. Overview of the multiscale simulation framework

We developed a multiscale simulation framework (**Figure 1**) to model cell-cell interactions in spatially organized tissues by coupling three complementary layers: a cell-resolved agent-based model (ABM), continuum partial differential equation (PDE) fields representing the tissue microenvironment, and intracellular Boolean gene-regulatory/state-transition networks. The framework is designed to bridge molecular, cellular, and tissue scales and to support integration of single-cell and spatial omics data. At the cellular level, each cell is represented as an individual agent with explicit position, phenotype, internal state, and interaction rules. At the tissue level, diffusible microenvironmental signals are represented as continuous fields on a 2D lattice. At the intracellular level, cell behaviors are controlled by compact Boolean state networks that translate local environmental and contact-dependent inputs into interpretable cell decisions such as activation, proliferation, cytotoxicity, suppression, and apoptosis.

**Figure 1.**
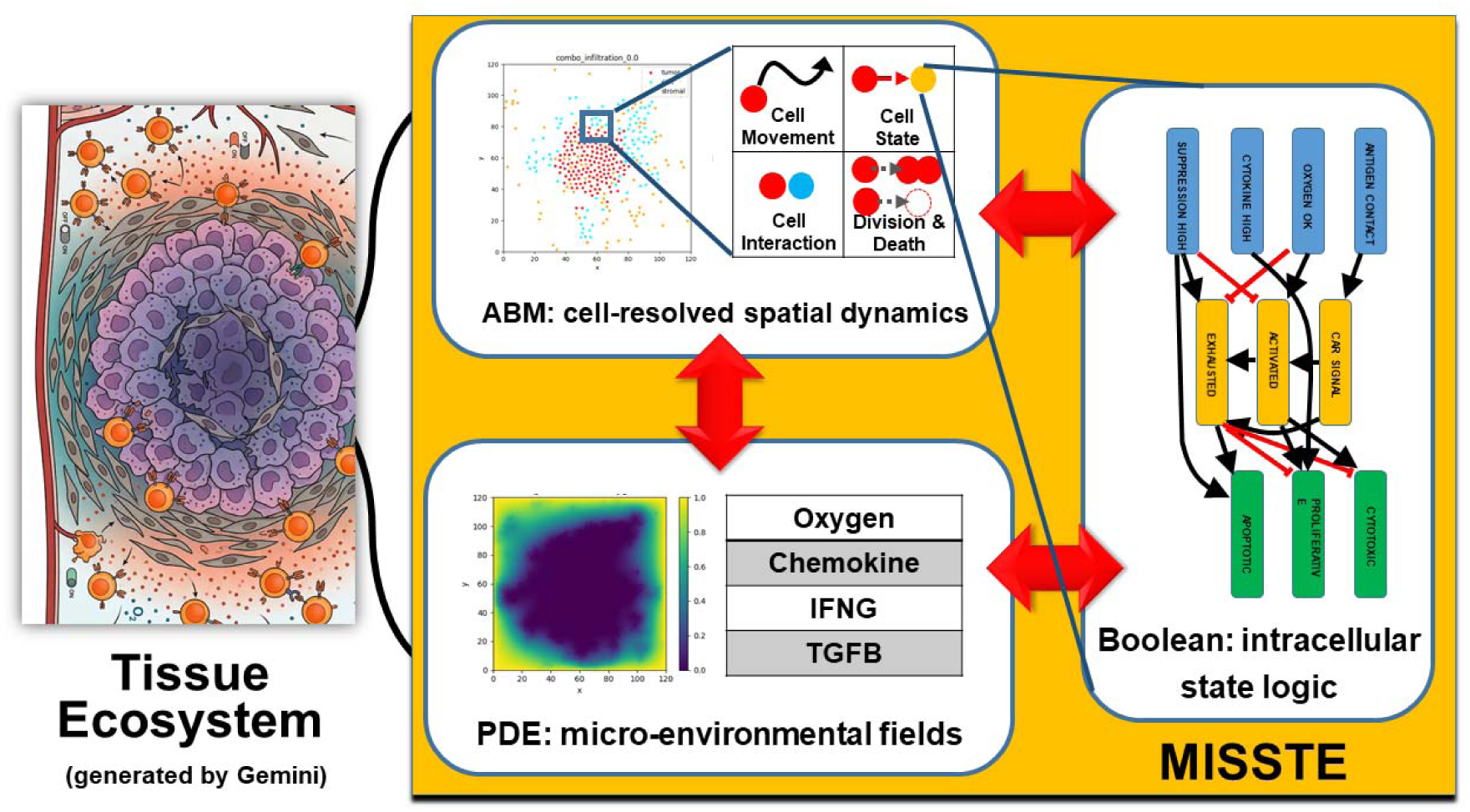
Overview of the MISSTE multiscale simulation framework. MISSTE integrates three coupled modeling layers to represent spatially organized tissue ecosystems: an agent-based model (ABM) for cell-resolved spatial dynamics, a partial differential equation (PDE) layer for diffusible microenvironmental fields, and Boolean intracellular state networks for cell-specific decision logic. The ABM captures cell movement, contact, proliferation, and death; the PDE layer represents continuous gradients such as oxygen, chemokine, inflammatory cytokines, and suppressive signals; and the Boolean layer maps local inputs into interpretable intracellular states that regulate cellular behavior. Red arrows indicate bidirectional coupling among layers. CAR-T therapy in a tumor microenvironment is used here as a proof-of-concept application.

The model is intentionally modular. Distinct biological systems can be instantiated by changing cell classes, state mappings, interface rules, and data-derived priors without altering the overall simulation architecture. In the present implementation, the framework was demonstrated in a multicellular system containing tumor, engineered immune, and stromal cells, but the underlying design is generalizable to other tissue ecosystems for which single-cell and spatial omics data are available.

### 2.2. Agent-based model of cell populations

#### Agent representation

Each cell was represented as an off-lattice agent with explicit spatial coordinates (x, y), cell identity, continuous low-dimensional state vector, intracellular Boolean state, and additional attributes such as antigen abundance or synthetic-cell identity. Agents belonged to discrete classes corresponding to biologically distinct populations, such as tumor, stromal, or engineered immune cells. Each agent was updated at every simulation step according to its local microenvironment, nearby cellular contacts, and internal logic state.

#### Cell movement

Agent motion was modeled in continuous space using a discrete-time position update. At each time step, the displacement of each cell was determined by a combination of stochastic motility, directed movement along local field gradients, and short-range repulsive interactions that prevented unrealistic overlap. For agent i, the update took the general form:

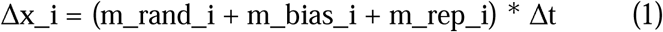

In above equation m_rand_i is a stochastic motility term scaled by cell-type-specific motility parameters, m_bias_i is a directed migration term derived from local gradients of diffusible fields or tissue-derived attraction fields, and m_rep_i is a short-range repulsion term computed from nearby cells.

For immune-like agents, directed migration could depend on chemokine gradients, tumor-density attraction, or contact-induced inward migration biases. For stromal and tumor cells, motion was typically weaker and governed primarily by stochastic motility and local repulsion, with optional biased responses to selected fields.

#### Cell-cell contact and neighborhood detection

Potential interactions between cells were determined by pairwise distance thresholds. Two cells were considered in contact if their Euclidean distance was less than a specified contact radius. These contact relationships were used to determine antigen engagement, local killing opportunities, and contact-dependent state updates. Additional neighbor lists within larger radii were used for local density estimation, crowding, and repulsion calculations.

#### Cell division and death

At each time step, agents were assigned division and death probabilities based on cell type, local environment, and intracellular state. These probabilities were interpreted as per-step stochastic event rates. When a division event occurred, a daughter cell was placed near the parent cell with inherited attributes and small perturbations to continuous internal state variables. Death events removed agents from the active population. This stochastic update scheme enabled local variability while preserving mechanistic control over state-dependent population dynamics.

### 2.3. Intracellular Boolean state networks

#### Rationale

To represent intracellular decision-making without introducing a large system of continuous intracellular equations, each agent was endowed with a compact Boolean state network [29–31]. This network receives local inputs from the environment and from direct cell-cell contact and updates a set of binary internal state nodes. These nodes then determine cell behaviors such as activation, proliferation, cytotoxicity, exhaustion, suppression, and apoptosis.

#### Input nodes

Boolean inputs were derived from local environmental measurements and contact relationships. Examples included antigen contact, adequate oxygen, high cytokine exposure, and high suppressive signaling. Thresholding continuous fields into Boolean inputs allowed environmental information from the PDE layer to be integrated into interpretable rule-based intracellular logic.

#### State nodes and update rules

Each cell type had its own Boolean network topology and update rules. For immune-like cells, representative nodes included CAR or receptor signaling, activation, cytotoxicity, proliferation, exhaustion, and apoptosis. For tumor cells, nodes could include stress, proliferation, and apoptosis. For stromal cells, nodes could include suppressive activation and remodeling-related states. These update rules were evaluated synchronously at each simulation step using local field values and contact-dependent inputs.

The Boolean formulation was used as a mechanistically interpretable abstraction rather than a full gene-regulatory model. It provides a compact representation of intracellular control logic while allowing direct coupling to cell behavior and tissue-scale dynamics. Detailed design of the cell-specific Boolean networks for the system can be found in the **Appendix** and **Figure 2**.

**Figure 2.**
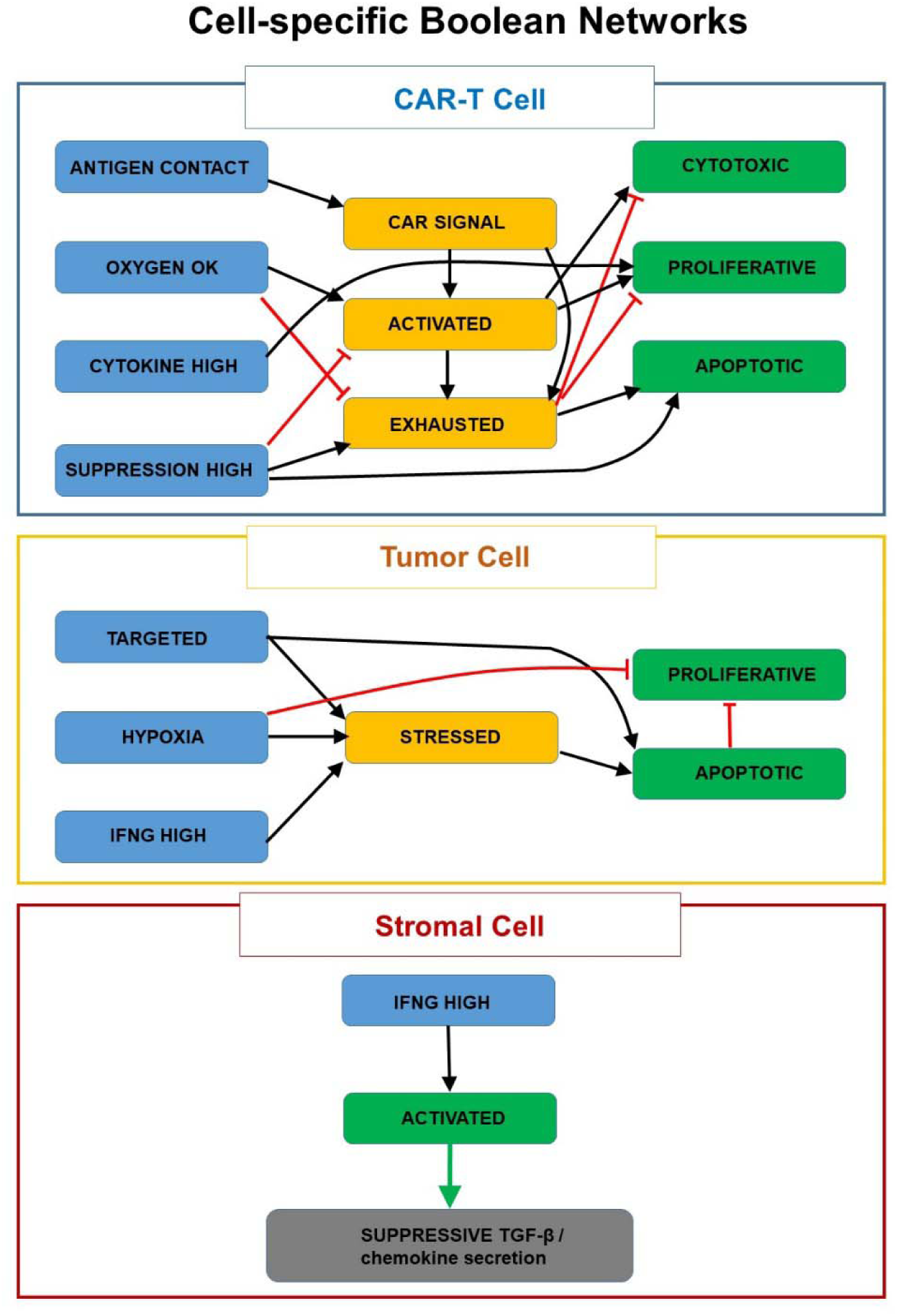
Cell-specific Boolean networks used in the multiscale framework. Schematic intracellular logic modules for CAR-T, tumor, and stromal cells. In the CAR-T network, antigen contact and local microenvironmental cues regulate CAR signaling, activation, proliferation, cytotoxicity, exhaustion, and apoptosis. In the tumor network, targeting, hypoxia, and inflammatory stress regulate stress, proliferation, and apoptosis. In the stromal network, inflammatory cues activate stromal cells and promote suppressive output, including TGF-beta and chemokine secretion. Black arrows denote activating relationships and red lines denote inhibitory relationships. These compact Boolean modules provide interpretable intracellular control logic that links local environmental input to cell behavior in the ABM-PDE simulation.

### 2.4. Continuum microenvironment modeled by PDEs

#### Field variables

The extracellular microenvironment was represented by a set of continuous fields defined on a 2D regular grid. In the current implementation, these fields included oxygen, chemokine, inflammatory cytokine-like signal, and suppressive signal. More generally, the framework can accommodate additional fields such as nutrients, metabolites, extracellular matrix modifiers, or region-specific molecular cues.

#### PDE formulation

Each field u_k(x,t) evolved according to a reaction-diffusion equation of the form:

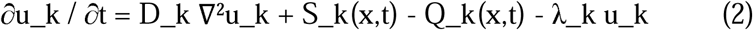

In above equation, D_k is the diffusion coefficient, S_k is the total production term from nearby cells, Q_k is the total consumption term, and λ_k is a decay coefficient. Production and consumption were determined dynamically from agent behaviors. For example, activated immune cells contributed inflammatory signals, stromal cells contributed suppressive factors, and multiple cell types consumed oxygen.

#### Numerical discretization

The PDE fields were updated on a regular grid using a finite-difference approximation to the Laplacian and explicit time stepping [32]. Spatial derivatives were discretized using a standard five-point stencil, and the fields were advanced using a forward Euler update. This scheme was chosen for transparency and ease of integration with the ABM layer. Boundary conditions could be specified per field; for example, oxygen fields could include boundary replenishment while other fields decayed or evolved without external source terms.

### 2.5. Coupling between ABM, PDE, and Boolean layers

The three model layers were coupled at every simulation step.

#### PDE to Boolean/ABM coupling

For each agent, local field values were sampled from the PDE grid at the agent’s position. These values were thresholded into Boolean inputs such as adequate oxygen, high cytokine exposure, or high suppressive signaling. In addition, local gradients of selected fields were used to generate directional movement biases. Thus, the PDE layer directly influenced both intracellular state updates and cell migration.

#### ABM to Boolean coupling

Direct cell-cell contact relationships generated Boolean input events such as antigen engagement or local targeting. These events were passed into the intracellular Boolean networks, allowing contact-dependent activation and killing logic to emerge from local spatial interactions.

#### Boolean to ABM coupling

After Boolean state updates, cell behaviors were recalculated. Internal states controlled whether a cell could proliferate, die, secrete diffusible factors, move more aggressively, or execute cytotoxic interactions. Thus, intracellular logic translated environmental and contact-dependent information into cell-level behaviors.

#### ABM to PDE coupling

Agent behaviors generated source and sink terms for the PDE fields. Cells secreted diffusible signals, consumed oxygen, and modified local field dynamics based on their current behavior states. This completed the feedback loop between cells and tissue environment.

Together, this yielded a closed multiscale update cycle in which tissue fields influence intracellular states, intracellular states influence cell behaviors, and cell behaviors reshape the tissue environment.

### 2.6. Parameter determination

The parameter values used in this study were not intended to represent directly fitted biophysical constants, but were chosen through a combination of biologically motivated initialization, mechanistic consistency, and iterative simulation-based refinement. Baseline values were first set to produce qualitatively reasonable tissue organization and cell-population behavior, including a central tumor mass, peripheral CAR-T localization, stromal expansion, and measurable but incomplete immune control. From this starting point, parameters governing contact, migration, killing, and suppressive interactions were varied within biologically plausible relative ranges to test how distinct mechanisms shape system-level outcomes. We then performed systematic parameter scans to identify which variables had the strongest influence on tumor burden, CAR-T persistence, and spatial coverage, and selected the final optimized and staged-strategy settings from these analyses as representative regimes that improved tumor control while remaining mechanistically interpretable. Thus, the current parameterization should be viewed as hypothesis-driven and exploratory, aimed at identifying dominant control principles in the model rather than establishing uniquely identifiable quantitative estimates. Detailed parameter values are listed in **Supplemental Materials**. In future work, these parameters can be more directly constrained and personalized using experimentally measured single-cell and spatial omics data.

### 2.7. Code and data availability

All relevant source codes can be found in the GitHub repository: https://github.com/wujah/MISSTE.

## 3. Results

### 3.1. Evaluation of Coordinated Parameter Optimization on Multicellular CAR-T Dynamics

We first compared two representative simulation scenarios to assess how coordinated tuning of interaction, migration, and infiltration parameters alters multicellular system behavior. The first scenario, referred to as the original baseline, represents the untuned reference model used prior to parameter refinement. In this setting, cells followed the default interaction and migration rules, with a baseline contact radius of 3.2 and baseline motility settings, and without additional enhancement of directed infiltration or CAR-T persistence. This scenario therefore reflects the initial model regime in which CAR-T cells are present at the tumor periphery but have limited effective contact coverage and minimal penetration into the tumor core. The second scenario, referred to as the optimized setting, incorporated the parameter combination identified through iterative exploration of the simulation design space. Relative to the original baseline, this optimized scenario simultaneously increased the effective range of cell-cell engagement, strengthened directed migration toward relevant gradients, enhanced CAR-T cytotoxic potential, and introduced a moderate combined infiltration program representing improved chemotactic responsiveness, stromal/ECM penetration, and contact-induced inward movement.

Thus, the key difference between the two scenarios is that the original baseline reflects a minimally tuned reference state with restricted immune-tumor interaction and poor deep tissue access, whereas the optimized setting represents a coordinated pro-infiltration, pro-contact, and pro-killing regime designed to improve tumor coverage, especially in spatially constrained regions. The comparison of these two scenarios was used to determine whether simultaneous optimization of interaction radius, migration bias, killing strength, and infiltration behavior could shift the system from a poorly penetrating baseline state toward improved CAR-T access and tumor control.

The simulation results are shown in **Figure 3**. For each scenario, five independent replicates were performed, with each trajectory running for 5,000 simulation steps. Mean trajectories with standard deviation showed consistent separation between the two regimes across tumor, CAR-T, stromal, and exhaustion dynamics. Relative to the original baseline, the optimized setting reduced tumor expansion throughout the simulation and yielded a lower final tumor burden (approximately 183 vs. 210 cells). The optimized regime also markedly improved CAR-T persistence: whereas CAR-T counts declined steadily in the baseline condition, they expanded and remained elevated under the optimized condition, reaching ∼58–60 cells at late times compared with ∼23 cells in baseline. Stromal accumulation was strongly suppressed in the optimized regime, with final counts remaining near ∼125–130 cells versus ∼445 cells in the baseline condition. In addition, the optimized setting showed a lower late-stage CAR-T exhaustion fraction, despite transient variability during the intermediate phase.

**Figure 3.**
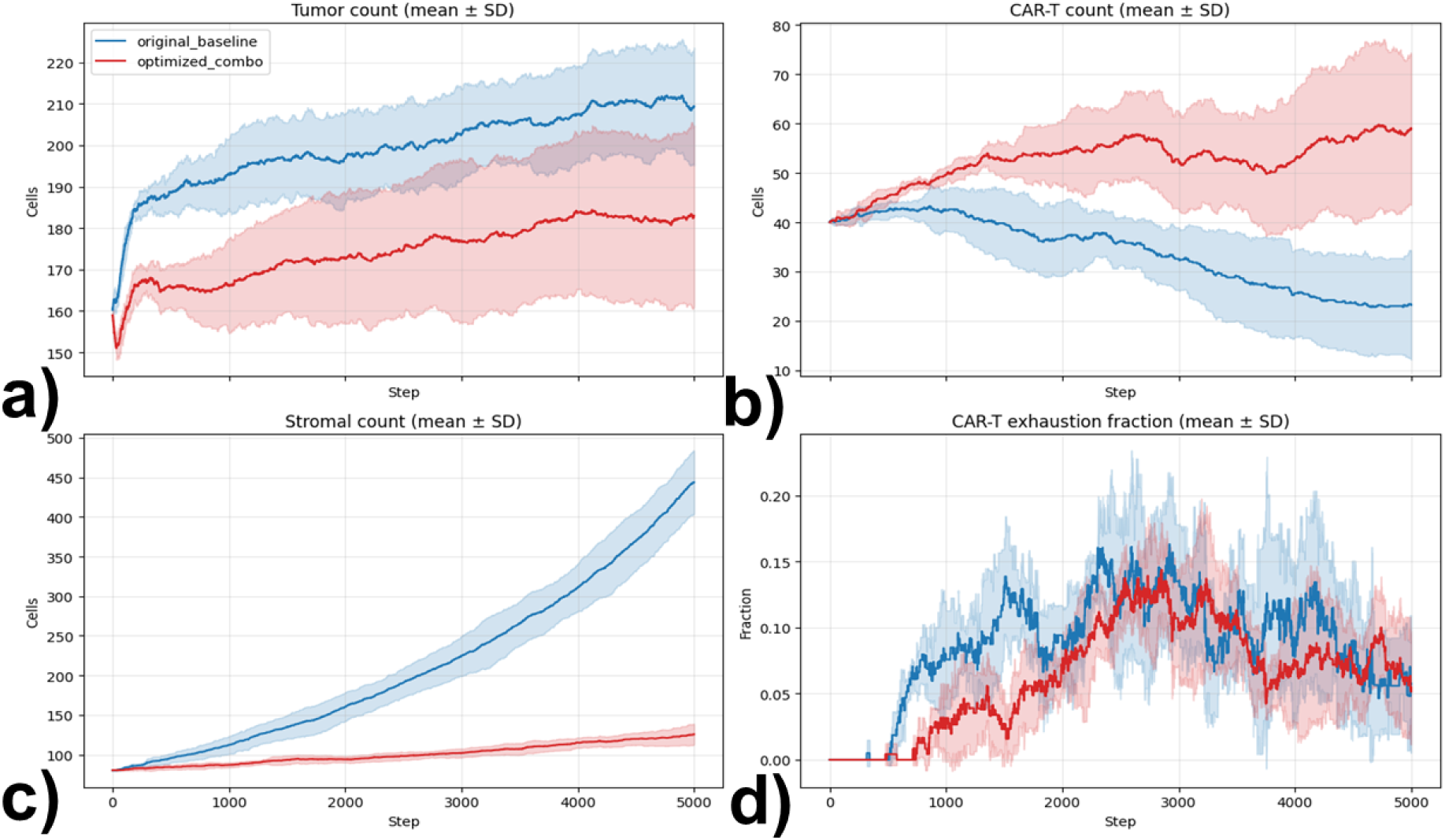
Comparison of baseline and optimized model trajectories across 5 replicates. Mean trajectories with standard deviation are shown for tumor count (a), CAR-T count (b), stromal count (c), and CAR-T exhaustion fraction (d) over 5000 simulation steps for the original baseline model (blue) and the optimized model (red). The optimized setting reduced tumor expansion, improved CAR-T persistence, limited stromal overgrowth, and lowered late-stage exhaustion relative to the baseline condition. Shaded bands represent variability across replicates.

Although replicate variability was present, especially for CAR-T abundance and exhaustion, the overall trends were robust across all 5 replicates. Together, these results indicate that coordinated increases in interaction range, directed migration, killing strength, and infiltration support shift the system from a stromal-dominated, CAR-T-depleting regime toward improved immune persistence and partial tumor control.

Spatial analysis revealed the microenvironmental differences between the baseline and optimized regimes (**Figure 4**). In the baseline condition, tumor cells remained concentrated in the center and were accompanied by extensive stromal accumulation, diffuse TGFB enrichment, and limited CAR-T presence near the tumor boundary. In contrast, the optimized setting preserved a larger CAR-T population around the tumor, markedly reduced stromal expansion, strengthened IFNG-associated immune activity, and restricted the spatial extent of suppressive TGFB signaling. Oxygen remained low in the tumor core in both conditions, indicating that hypoxia persisted as a central barrier despite optimization. Together, these results suggest that the optimized regime improves tumor control primarily by reducing stromal suppression and enhancing local immune persistence, rather than by fully overcoming core protection.

**Figure 4.**
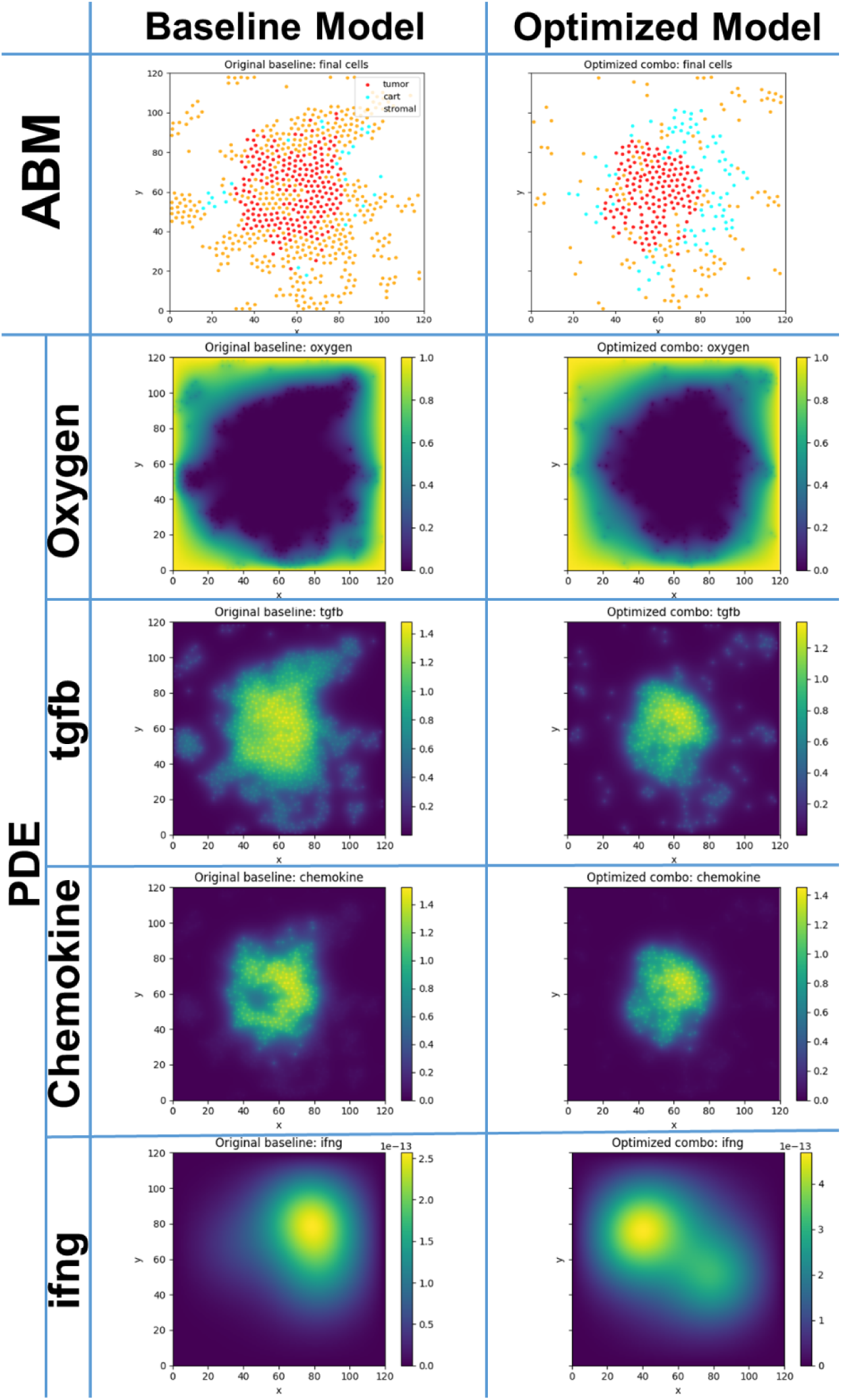
Spatial organization and microenvironmental field remodeling in the baseline and optimized models. Final spatial distributions of cells and PDE fields comparing the baseline and optimized conditions. The top row shows the final ABM cell positions for tumor, CAR-T, and stromal populations. The lower rows show representative PDE fields, including oxygen, TGF-beta, chemokine, and IFNG. Relative to the baseline, the optimized model preserves more CAR-T cells near the tumor, reduces stromal dominance, and alters the spatial distribution of suppressive and inflammatory signals. Together, these maps illustrate how optimization reshapes both cell organization and the surrounding microenvironment.

### 3.2. Sensitivity Analysis and Parameter Scans of Mechanistic Components Shaping Tumor-Immune Interactions

To identify which mechanistic components most strongly shape system behavior, we further performed one-factor-at-a-time parameter scans around the optimized backbone. We focused on four parameters chosen to represent distinct biological levers of CAR-T function in solid tumors: 1) contact radius, which approximates the effective spatial range over which CAR-T cells can engage and act on nearby tumor cells; 2) chemotaxis strength, which controls directed migration toward chemoattractive signals and reflects trafficking efficiency; 3) CAR-T kill base, which represents intrinsic cytotoxic potency once productive contact is established; and 4) core-attraction strength, which models an additional directed bias toward the tumor-dense interior and serves as a simplified proxy for mechanisms that promote deeper penetration into the tumor core. These four parameters were selected because they map onto biologically meaningful and experimentally tunable processes, including target engagement, migration, killing, and tissue penetration, and also because earlier analyses suggested that all these four factors could plausibly influence tumor control. In practice, each parameter was scanned over a broad range while the other three were held fixed. Every condition was run with 5 independent replicates, yielding 140 total simulations across the four scan panels.

The scans, as shown in **Figure 5**, revealed a clear hierarchy of importance. Among the four parameters, contact radius had the strongest effect on final tumor burden. Increasing contact radius produced a pronounced and approximately monotonic decrease in final tumor size, indicating that the effective interaction range between CAR-T and tumor cells is a dominant determinant of control in this model. This result is biologically consistent with the idea that even modest improvements in how frequently and how deeply CAR-T cells establish productive contact can substantially improve tumor clearance.

**Figure 5.**
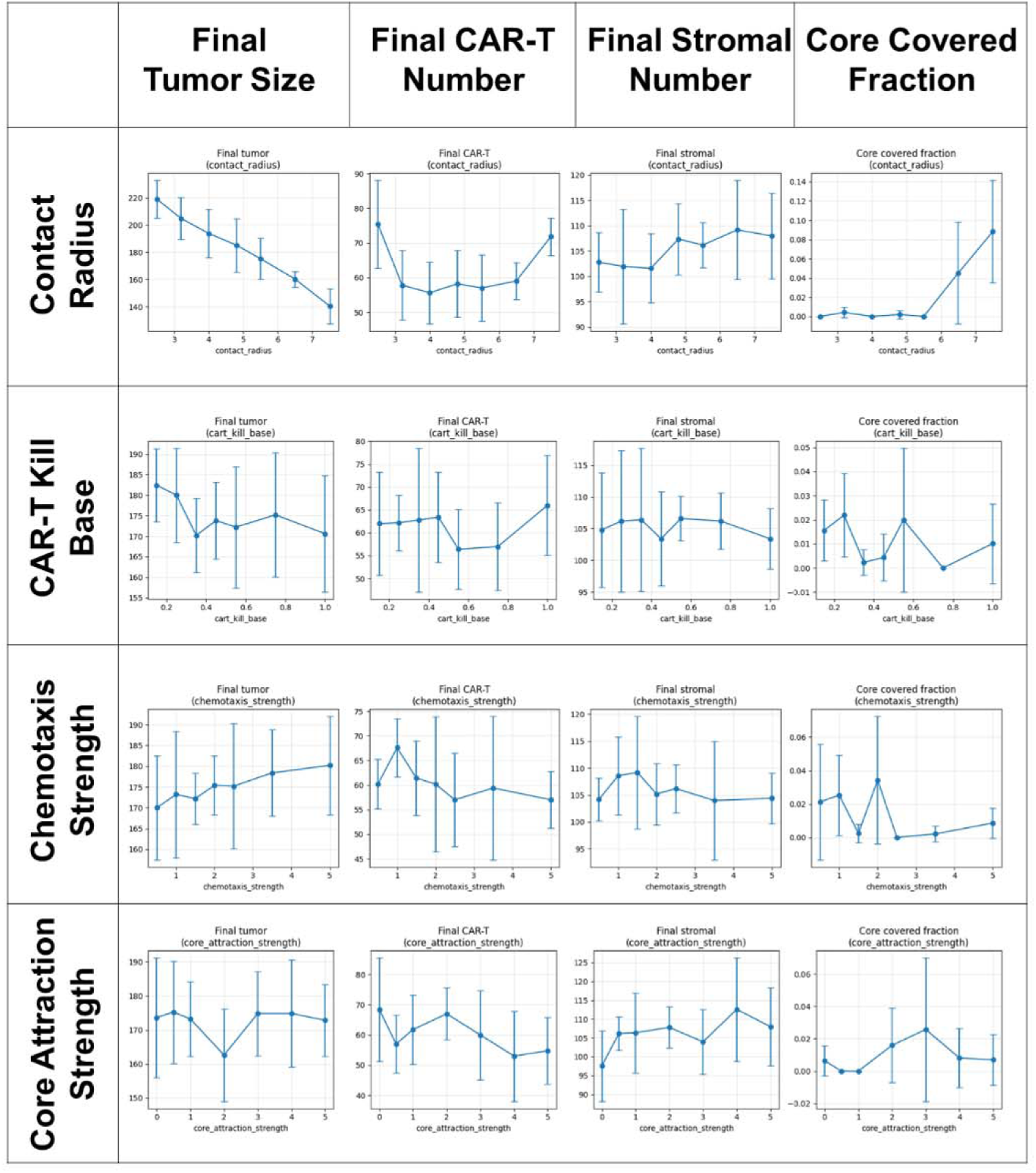
One-factor-at-a-time parameter scans identify dominant control variables in the model. Parameter scans were performed by varying contact radius, CAR-T kill base, chemotaxis strength, and core-attraction strength while holding the remaining parameters fixed at the optimized backbone. For each parameter value, final tumor size, final CAR-T number, final stromal number, and tumor core covered fraction were quantified across 5 replicates. Contact radius produced the strongest monotonic reduction in final tumor burden and the clearest increase in core coverage, whereas CAR-T kill base had a weaker effect, chemotaxis strength showed little benefit, and core-attraction strength had minimal impact on final tumor size. Error bars represent standard deviation across replicates.

By comparison, CAR-T kill base showed only a weak decreasing effect on final tumor burden. Increasing intrinsic killing potency modestly improved tumor control, but the effect size was far smaller than that observed for contact radius. This suggests that cytotoxic strength alone is not the principal bottleneck once CAR-T cells remain spatially restricted. In other words, making CAR-T cells more potent has limited benefit if they still fail to reach enough tumor cells.

In contrast, increasing chemotaxis strength produced a weak upward trend in final tumor burden rather than a reduction. Although stronger chemotaxis might be expected to improve immune recruitment, in this model it did not translate into improved tumor control and may instead have promoted more peripheral accumulation or inefficient clustering without sufficient deep engagement. Finally, core-attraction strength had little consistent effect on final tumor size across the range tested. While this parameter was intended to represent deeper inward bias toward the tumor interior, the scan indicates that this mechanism alone is insufficient to substantially alter overall tumor outcome.

Taken together, these results show that the model is most sensitive to parameters governing effective spatial engagement, rather than migration strength, inward bias, or killing potency in isolation. The strong dependence on contact radius, combined with the comparatively weak effects of kill strength and the negligible effect of core attraction, supports the conclusion that productive immune-tumor contact is the dominant bottleneck in this system. This finding provided the mechanistic basis for the subsequent design of staged intervention strategies aimed first at improving access and engagement, and only then at enhancing cytotoxic function and limiting late-stage dysfunction.

### 3.3. Development and Evaluation of Temporally Ordered Therapeutic Interventions

Following parameter scans, we next asked whether the model could be used not only to identify important control parameters, but also to design time-dependent intervention strategies for stronger tumor reduction. The rationale for this final analysis came directly from the earlier results. First, the baseline and optimized comparisons indicated that improved CAR-T persistence and reduced stromal suppression were beneficial, but were not by themselves sufficient to eliminate the tumor. Second, the parameter scans showed that effective spatial engagement, particularly contact range, had a much larger impact on tumor control than simply increasing killing potency, whereas stronger chemotaxis alone or inward core attraction alone had limited benefit. Together, these findings suggested that the dominant bottlenecks are likely to arise in sequence: CAR-T cells must first gain effective access to the tumor, then exploit that access through a period of stronger cytotoxic function, and finally avoid late functional decline in the suppressive microenvironment. On this basis, we specifically designed four strategies with increasing mechanistic complexity: 1) S1, a constant optimized reference condition using the same setting as in the first part as control; 2) S2, which transiently enhances early access and contact; 3) S3, which adds a subsequent period of elevated cytotoxicity after infiltration has been established; and 4) S4, which further introduces a late maintenance phase aimed at limiting exhaustion and suppressive stromal influence. Every condition was run with 5 independent replicates. This staged design was intended to test the broader hypothesis that effective tumor control may require temporally ordered intervention, rather than a single static enhancement applied throughout the simulation.

Figure 6 shows that all three staged strategies (S2-S4) outperformed the static optimized reference in terms of final tumor burden (Figure 6a and **6e**). The trajectory plots indicate that this benefit emerged early and was then maintained. Under S1, tumor burden increased steadily after the initial transient and remained clearly separated from the other strategies throughout most of the simulation. In contrast, S2, S3, and S4 all drove an early drop in tumor burden followed by a lower plateau. This suggests that early improvement in tumor access is a key determinant of the eventual outcome and that the static optimized regime remains limited by insufficient effective engagement despite otherwise favorable parameters.

**Figure 6.**
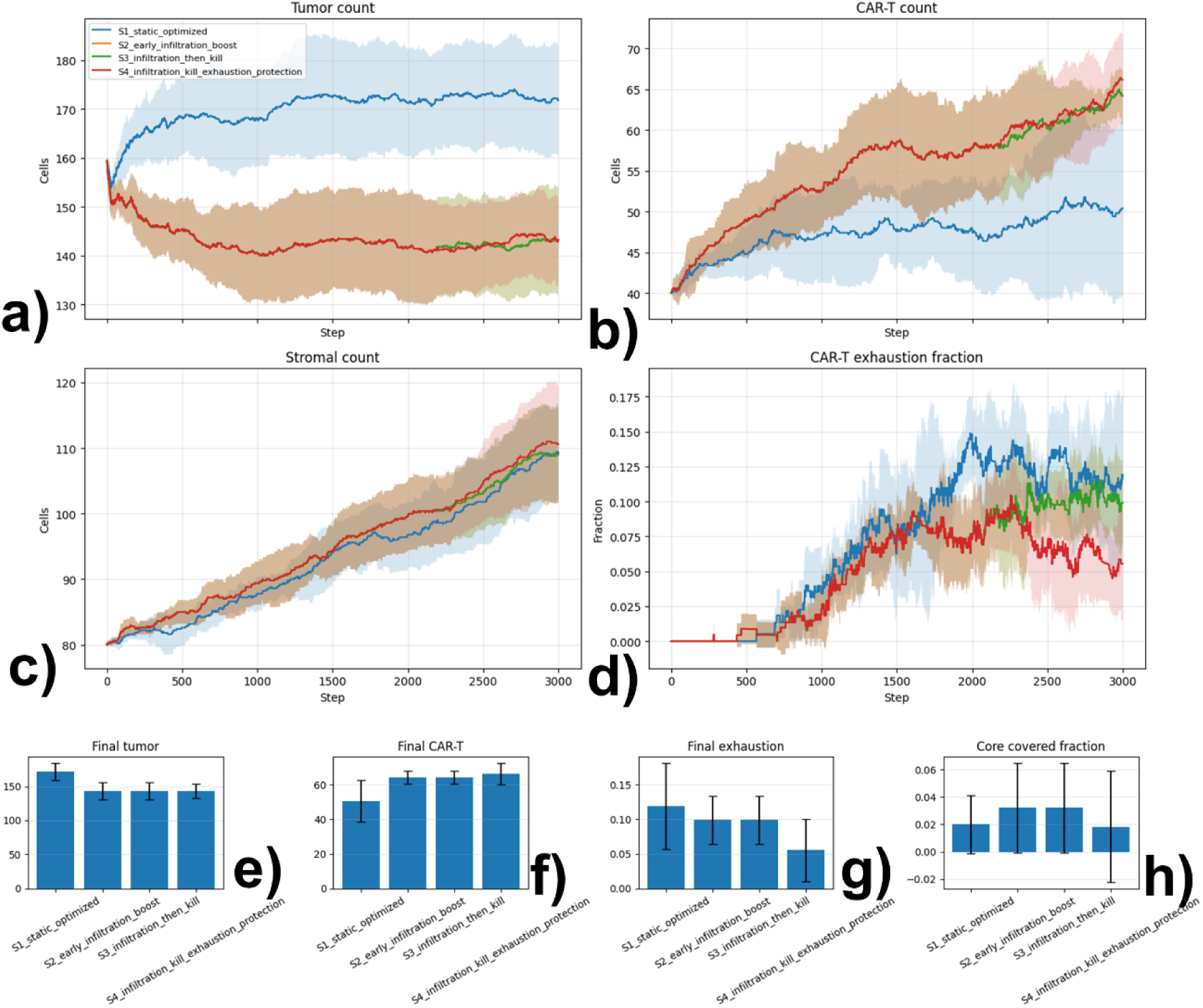
Sequential intervention strategies improve tumor control relative to the static optimized regime. Four dynamic treatment strategies were compared: S1, static optimized control; S2, early infiltration boost; S3, early infiltration followed by a kill boost; and S4, infiltration followed by kill boost and late exhaustion protection. Panels (a–d) show mean trajectories with standard deviation for tumor count, CAR-T count, stromal count, and CAR-T exhaustion fraction across 5 replicates. Panels (e–h) summarize final tumor burden, final CAR-T abundance, final exhaustion fraction, and tumor core covered fraction. All staged strategies reduced final tumor burden relative to the static optimized condition, while S4 produced the lowest final exhaustion fraction, consistent with improved late-stage maintenance of CAR-T function.

The dynamic strategies also increased CAR-T persistence (Figure 6b and **6f**). Final CAR-T abundance rose from approximately 50 cells under S1 to roughly 64–66 cells under S2–S4. This increase was visible throughout the trajectories, with staged strategies supporting a larger and more sustained CAR-T population over time. Stromal counts were comparatively similar across all four conditions by the end of the simulation (Figure 6c), indicating that the principal separation among strategies was not driven by major changes in final stromal abundance, but rather by differences in CAR-T persistence and functional state.

Among the staged strategies, the differences in final tumor burden were relatively small, indicating that early infiltration enhancement alone already captured most of the benefit. Adding a subsequent kill-boost phase (S3) did not substantially reduce tumor burden beyond S2, and the full three-stage strategy (S4) produced tumor control comparable to the other staged schedules. This result is notable because it suggests that once spatial access is improved, additional escalation of cytotoxicity yields only limited extra benefit in this model.

However, the late protection phase in S4 had a clear effect on CAR-T functional state. S4 achieved the lowest final exhaustion fraction, approximately 0.055, compared with values near 0.10–0.12 in the other conditions (Figure 6d and **6g**). The exhaustion trajectories further showed that S4 diverged downward at late times, consistent with the intended maintenance/protection phase. Thus, while the added exhaustion-protection stage did not dramatically outperform the other staged strategies in final tumor count, it improved the quality of the CAR-T response by preserving a less exhausted population.

Taken together, these results show that sequential, mechanism-guided intervention schedules outperform a static optimized regime, and that the major gain comes from addressing the bottlenecks in the correct order. Early enhancement of infiltration and engagement appears to be the most important step for reducing tumor burden, whereas later kill boosting and exhaustion protection mainly shape the persistence and functional quality of the CAR-T population. This supports the broader conclusion that in multiscale tissue ecosystems, effective intervention may require temporally ordered control of access, effector function, and maintenance, rather than uniform enhancement of all properties throughout treatment. Clinically, the staged strategies can be interpreted as temporally ordered combination regimens in which early interventions improve tumor access, intermediate interventions transiently enhance cytotoxic function after engagement has been established, and late interventions preserve CAR-T fitness by limiting exhaustion and suppressive microenvironmental pressure.

## 4. Discussions

Tissue ecosystems are organized communities of interacting cell types embedded within spatially structured biochemical and physical microenvironments. Their behavior emerges not only from the intrinsic programs of individual cells, but also from local cell-cell contact, competition, cooperation, and diffusible signals that vary across space and time. Understanding these systems that operate across molecular, cellular, and tissue scales requires computational frameworks that can unify intracellular decision-making, spatial cell-cell interactions, and continuous microenvironmental regulation within a single mechanistic model. In this study, we developed MISSTE as a modular multiscale framework that integrates Boolean intracellular logic, agent-based cellular dynamics, and PDE-based tissue fields, and applied it to CAR-T therapy in solid tumors as a proof-of-concept case study. Using this system, we showed that the framework can reproduce emergent behaviors that are difficult to infer from any single scale alone, including the coupled effects of immune-tumor contact, stromal suppression, cytokine field remodeling, and functional exhaustion.

A recent experimental study provides a useful biological parallel to our modeling results. Pang et al. engineered FAP/IL-15 CAR-T cells to simultaneously deplete FAP-positive cancer-associated fibroblasts and enhance CAR-T survival, proliferation, and memory-like function through endogenous IL-15 secretion [33]. Their rationale was that solid-tumor failure arises not only from insufficient cytotoxicity, but also from stromal/ECM barriers, impaired trafficking, limited intratumoral persistence, and rapid exhaustion within the immunosuppressive tumor microenvironment. This is highly consistent with our simulations, in which improved tumor control was not achieved by stronger killing alone, but instead emerged when stromal restriction was relieved and CAR-T persistence/infiltration were jointly improved. Our model extends these experimental observations by offering a spatially explicit mechanistic interpretation: beyond stromal suppression and T-cell fitness, a key limiting factor may be incomplete coverage of the tumor core. In this view, stromal depletion and persistence-promoting programs such as IL-15 do not simply increase overall CAR-T activity; they may act by enabling deeper tumor access, sustaining local effector function, and preventing peripheral trapping. Thus, our framework is consistent with the experimental findings of Pang et al., while adding the new insight that tumor-core access and durable spatial coverage may be the proximate determinants separating partial control from complete collapse in solid-tumor CAR-T therapy.

A natural next step for MISSTE is the incorporation of experimentally measured single-cell and spatial omics data to replace stylized initial conditions and further constrain model structure. In such a data-driven extension, single-cell profiles could be used to define cell classes, infer low-dimensional intracellular state variables, and initialize Boolean logic modules or state-transition biases, while spatial omics data could provide cell coordinates, tissue domains, neighborhood structure, and region-specific microenvironmental priors. These measurements could be mapped onto simulation agents, local interaction rules, and PDE field initializations, allowing the model to begin from experimentally observed tissue organization rather than synthetic configurations. In this setting, MISSTE would serve not only as a forward simulator but also as a platform for integrating spatially resolved molecular data with mechanistic inference, enabling more realistic reconstruction of tissue ecosystems and more quantitative hypothesis testing across conditions, perturbations, and patient-specific contexts.

Beyond CAR-T therapy, MISSTE is designed as a general framework for studying multicellular systems in which intracellular decision logic, spatial cell-cell interactions, and diffusible tissue signals jointly determine system behavior. The same architecture could be applied to other engineered immune therapies, including TCR-T cells [34–36], NK-cell therapies [37–39], macrophage-based therapies [40–42], and combinatorial cell products in which activation, trafficking, persistence, and dysfunction emerge from local tissue context. It is also readily extensible to non-therapy settings, such as tumor–stroma–immune crosstalk, wound repair, fibrotic remodeling, developmental patterning, host–pathogen interactions, and regenerative medicine, where discrete cell behaviors are shaped by both local contact rules and continuous morphogen or cytokine fields. More broadly, because the framework separates the PDE layer, the agent rules, and the Boolean intracellular modules into interchangeable components, it can be re-parameterized for diverse tissue ecosystems using spatial omics, single-cell profiling, or perturbation data without changing the core simulation engine. In this sense, the CAR-T system studied here not only serves as a biologically important application, but also as a proof-of-concept demonstrating how a unified multiscale formalism can be used to interrogate a wide class of spatially organized biological systems.

## Supporting information

Parameter used in the simulations

## 5. Appendix: Cell-specific Boolean networks

To represent intracellular decision-making in a compact and interpretable form, each simulated cell carries a cell-type-specific Boolean state network. These networks map local environmental conditions and contact-dependent cues to discrete intracellular states that regulate behavior at the agent level. Boolean nodes take binary values and are updated at each simulation step from thresholded field inputs and local cell-cell interactions. The resulting state configuration determines whether a cell becomes activated, proliferative, cytotoxic, suppressive, stressed, or apoptotic.

The networks are shown in **Figure 2**. The figure is structured as directed logic graphs flowing from left to right. Blue nodes represent thresholded environmental inputs received from the tissue (PDE) or contact (ABM) layers. Orange nodes represent internal computed states. Green nodes represent the final behavioral outputs executed by the cell. Gray nodes indicate constant, invariant states. Black arrows (→) denote activating relationships or necessary conditions (AND/OR logic), while red flat-ended lines (–|) denote inhibitory relationships (NOT logic).

### CAR-T cell Boolean network

The CAR-T Boolean network encodes the transition from target recognition to functional activation or dysfunction. Input nodes include ANTIGEN_CONTACT, OXYGEN_OK, CYTOKINE_HIGH, and SUPPRESSION_HIGH, which are derived from local contact status and sampled microenvironmental fields. These inputs regulate the internal state nodes CAR_SIGNAL, ACTIVATED, CYTOTOXIC, PROLIFERATIVE, EXHAUSTED, and APOPTOTIC.

Antigen engagement activates CAR_SIGNAL, which in turn promotes ACTIVATED. Activation is further supported by adequate oxygen and is inhibited by strong suppressive signaling. The ACTIVATED state promotes CYTOTOXIC behavior and, together with high cytokine exposure, can induce the PROLIFERATIVE state. In parallel, persistent target engagement under suppressive conditions can induce EXHAUSTED, which suppresses both cytotoxicity and proliferation. When exhaustion is combined with strong suppression, the APOPTOTIC node is activated, leading to increased death probability. In this way, the CAR-T network captures the balance between productive target engagement and dysfunctional overstimulation.

### Tumor cell Boolean network

The tumor Boolean network represents the balance between growth, stress, and death under immune and microenvironmental pressure. Representative input nodes include OXYGEN_LOW, IFNG_HIGH, IMMUNE_CONTACT, and NUTRIENT_OK. These regulate the internal state nodes STRESS, SURVIVAL, PROLIFERATIVE, and APOPTOTIC.

Hypoxia, inflammatory signaling, and immune contact promote STRESS, while nutrient sufficiency promotes SURVIVAL and supports PROLIFERATIVE behavior. Stress inhibits proliferation and can activate apoptosis when sufficiently strong. Immune contact can also reduce tumor survival signaling. This logic allows tumor cells to switch between growth-favoring and damage-responsive states depending on local immune pressure and metabolic context.

### Stromal cell Boolean network

The stromal Boolean network encodes activation of matrix-remodeling and suppressive stromal programs. Input nodes include IFNG_HIGH, DAMAGE_SIGNAL, TGFB_HIGH, and HYPOXIA, which reflect inflammatory, injury-related, suppressive, and metabolic signals in the local environment. These inputs regulate internal state nodes such as ACTIVATED_STROMA, SUPPRESSIVE, REMODELING, and QUIESCENT.

Damage-associated and inflammatory inputs promote ACTIVATED_STROMA, which can then drive both SUPPRESSIVE and REMODELING programs. High TGF-beta further reinforces the suppressive state, while hypoxia can promote remodeling-related behavior. Activated or suppressive stromal states inhibit quiescence, thereby shifting stromal cells toward active participation in tissue restructuring and immune exclusion.

### Coupling of Boolean states to agent behavior

The Boolean state of each cell is used to modulate behavior in the agent-based layer. For CAR-T cells, Boolean outputs alter migration bias, killing probability, proliferation, and death. For tumor cells, Boolean outputs determine proliferative capacity and apoptosis susceptibility. For stromal cells, Boolean outputs control suppressive factor production, matrix-remodeling behavior, and persistence. Ths formulation provides a mechanistically interpretable intracellular layer that couples local microenvironmental sensing to emergent multicellular dynamics.

## Acknowledgement

This work was supported by the National Institutes of Health under Grant Numbers R01GM122804, and the United States – Israel Binational Science Foundation Project Number: 2023336. The work is also partially supported by a start-up grant from Albert Einstein College of Medicine. Computational support was provided by Albert Einstein College of Medicine High Performance Computing Center. Finally, we also greatly appreciate the general support from the Einstein 2030 Seed Fund.

## Author Contributions

Z.S., S.Y. and Y.W. designed research; Z.S. and Y.W. performed research; Z.S., and Y.W. analyzed data; Z.S. and Y.W. wrote the paper.

